# The intracellular C-terminus confers compartment-specific targeting of voltage-gated Ca^2+^ channels

**DOI:** 10.1101/2023.12.23.573183

**Authors:** Morven Chin, Pascal S. Kaeser

## Abstract

To achieve the functional polarization that underlies brain computation, neurons sort protein material into distinct compartments. Ion channel composition, for example, differs between axons and dendrites, but the molecular determinants for their polarized trafficking remain obscure. Here, we identify the mechanisms that target voltage-gated Ca^2+^ channels (Ca_V_s) to distinct subcellular compartments. In hippocampal neurons, Ca_V_2s trigger neurotransmitter release at the presynaptic active zone, and Ca_V_1s localize somatodendritically. After knockout of all three Ca_V_2s, expression of Ca_V_2.1, but not of Ca_V_1.3, restores neurotransmitter release. Chimeric Ca_V_1.3 channels with Ca_V_2.1 intracellular C-termini localize to the active zone, mediate synaptic vesicle exocytosis, and render release fully sensitive to blockade of Ca_V_1 channels. This dominant targeting function of the Ca_V_2.1 C-terminus requires an EF hand in its proximal segment, and replacement of the Ca_V_2.1 C-terminus with that of Ca_V_1.3 abolishes Ca_V_2.1 active zone localization. We conclude that the intracellular C-termini mediate compartment-specific Ca_V_ targeting.

## Introduction

Neurons are polarized cells with a defined signaling directionality from dendrites to soma to axon ^1^. To achieve this morphological and functional polarization, neurons sort protein material into specific subcellular compartments ^2,3^. Voltage-gated Ca^2+^ channels (Ca_V_s), which couple electrical activity to changes in intracellular Ca^2+^ signaling, are a prototypical example of sorting specificity. They are a large protein family, and individual members localize to distinct subcellular domains in the dendrites, the soma and the axon ^4,5^. However, Ca_V_ subtypes exhibit limited differences in their sequences, and the molecular determinants that target Ca_V_s to specific subcellular compartments remain elusive.

Ca_V_s are defined by their pore-forming Ca_V_α1 subunit, and their expression, trafficking and function are modulated by Ca_V_β subunits and Ca_V_α2δ proteins ^4–7^. Vertebrate Ca_V_α1 subunits are encoded by ten genes classified into Ca_V_1 (Ca_V_1.1-1.4, L-type), Ca_V_2 (Ca_V_2.1-2.3, P/Q-, N- and R-type), and Ca_V_3 (Ca_V_3.1-3.3, T-type) channels. Most Ca_V_s are abundantly co-expressed in central neurons. Ca_V_1.2 and Ca_V_1.3 have important roles in the somatodendritic compartment. There, Ca^2+^ influx through Ca_V_1 channels activates effectors to induce gene transcription ^8–11^ and modulates neuronal firing directly and through Ca^2+^-activated K^+^ channels ^12–16^. In presynaptic nerve terminals, Ca_V_2.1 (P/Q-type) and Ca_V_2.2 (N-type) are the primary Ca^2+^ sources for synaptic vesicle release ^17–20^. They are recruited to a specialized release apparatus, the active zone, where they are tethered near fusion-competent vesicles ^21–25^. This organization couples action potential-induced Ca^2+^ entry to vesicular release sites for the rapid and robust triggering of neurotransmitter exocytosis. Overall, Ca_V_s contribute to diverse cellular processes, and their functions are directly tied to their subcellular localization.

The mechanisms that distinguish Ca_V_1s from Ca_V_2s and sort them into the somatodendritic and axonal compartments, respectively, remain unclear. Starting from their primary site of synthesis in the soma, Ca_V_s likely undergo a series of interactions that target each subtype to its respective subcellular domain ^2,26^. However, Ca_V_s are highly similar in structure ^5,27,28^, and notable overlap exists within the Ca_V_1 and Ca_V_2 interactome. For example, interactions with Ca_V_β, Ca_V_α2δ, and calmodulin have been implicated in Ca_V_ trafficking ^29–34^, but these proteins interact indiscriminately with Ca_V_1s and Ca_V_2s and are thus unlikely to encode specific sorting information. The intracellular Ca_V_ C-termini might mediate targeting specificity. Ca_V_ C-termini include a proximal segment with two EF hands and an IQ motif, and a distal segment containing binding sites for scaffolding proteins (Figs. S1A+B). The Ca_V_2 C-terminus binds to the PDZ domain of the active zone protein RIM, and it contains a proline-rich sequence (which is also present in Ca_V_1s) that binds to RIM-BP ^24,35,36^. Together, these interactions help tether Ca_V_2s to the presynaptic active zone ^20,24,37–42^. Analogous sequences in Ca_V_1.3 bind to the postsynaptic scaffold Shank, and overall, Ca_V_1 C-termini support cell surface expression and the assembly of Ca_V_1 into dendritic clusters ^43,44^. An additional poly-arginine motif specific to Ca_V_2.1 may also contribute to its localization ^20,45^. Sequences outside the C-terminus could also be involved. For example, binding of the Ca_V_2 cytoplasmic II-III loop to SNARE proteins ^46–48^ and Ca_V_ interactions with material in the synaptic cleft may mediate anchoring at presynaptic sites ^49,50^. Taken together, multiple interactions have been implicated in Ca_V_ trafficking and targeting, but how these interactions direct Ca_V_1s and Ca_V_2s to opposing compartments has remained unclear.

Here, we found that the Ca_V_ C-termini are the primary determinants of channel localization in hippocampal neurons. Swapping the Ca_V_2.1 C-terminus onto Ca_V_1.3 targets the channel to the presynaptic active zone in Ca_V_2 knockout neurons. This chimeric Ca_V_1.3 channel mediates Ca^2+^ entry for neurotransmitter release and renders synaptic vesicle exocytosis sensitive to L-type Ca_V_ blockers. In contrast, the inverse swap prevents active zone localization of Ca_V_2.1. Within the Ca_V_2.1 proximal C-terminus, an EF hand is required for presynaptic targeting, and its removal leads to loss of Ca_V_2.1 from the active zone. We conclude that the C-terminus specifies Ca_V_ localization, and we identify the EF hand as an essential trafficking motif.

## Results

### Lentivirally expressed Ca_V_2.1, but not Ca_V_1.3, localizes to active zones and mediates neurotransmitter release after Ca_V_2 ablation

To determine the Ca_V_ sequences important for active zone localization, we expressed various Ca_V_s using lentiviruses in cultured hippocampal neurons that lack Ca_V_2.1, Ca_V_2.2 and Ca_V_2.3. Specifically, we transduced neurons that contain “floxed” conditional knockout alleles for these three channels (Fig. 1A) with lentiviruses that express cre recombinase under a synapsin promoter to generate Ca_V_2 cTKO neurons ^20^. Control neurons (Ca_V_2 control) were identical except for transduction by a lentivirus expressing a truncated, recombination-deficient version of cre. In addition, we transduced Ca_V_2 cTKO neurons with either a lentivirus expressing HA-tagged Ca_V_2.1 or with a lentivirus expressing HA-tagged Ca_V_1.3. The tags were inserted near the Ca_V_ N-terminus in a position shown previously to not interfere with the expression (Figs. 1B, S1A-1E), targeting and function of Ca_V_2.1 ^20,51^. We then used stimulated emission depletion (STED) microscopy (Fig. 1C-H), confocal microscopy (Fig. S1F-I), and electrophysiology (Fig. 1I-L) to assess Ca_V_ localization and synaptic transmission.

**Figure 1.**
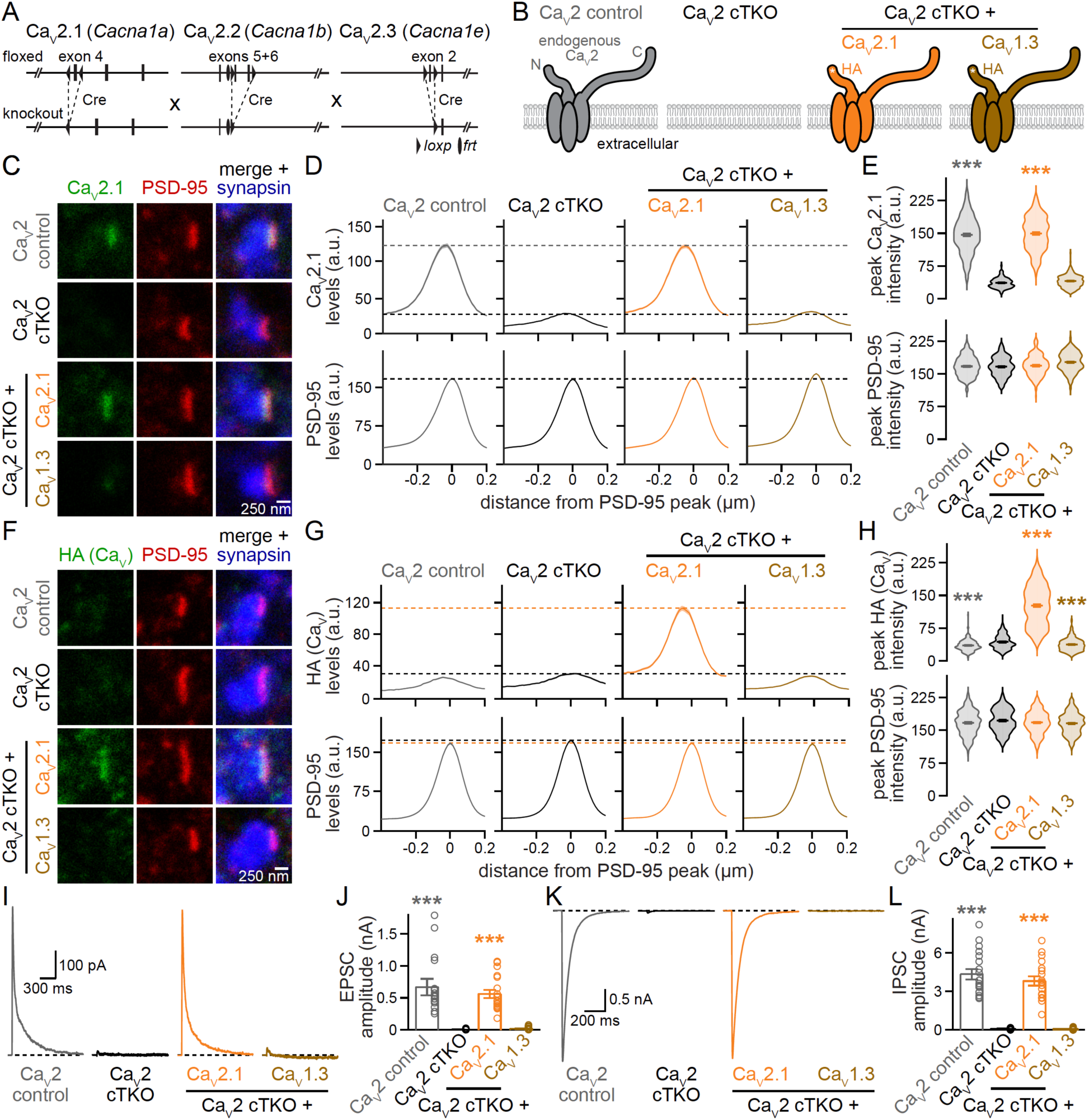
Lentivirally expressed Ca_V_2.1, but not Ca_V_1.3, localizes to active zones and restores synaptic transmission in Ca_V_2 triple knockout neurons. (A) Strategy for Ca_V_2 triple knockout in cultured hippocampal neurons as described before ^20^. Transduction of neurons from triple-floxed mice with a lentivirus expressing Cre recombinase produced Ca_V_2 cTKO neurons, Ca_V_2 control neurons were identical except for the expression of a truncated, recombination-deficient Cre. (B) Schematic of the conditions for comparison (schematics similar to ^20^); HA-tagged (HA) Ca_V_s were expressed by lentiviral transduction. (C-E) Representative images (C) and summary plots of intensity profiles (D) and peak levels (E) of Ca_V_2.1 and PSD-95 at synapses in side-view, levels are shown in arbitrary units (a.u.). Neurons were stained with antibodies against Ca_V_2.1 (analyzed by STED microscopy), PSD-95 (STED), and synapsin (confocal). Dashed lines in D denote levels in Ca_V_2 cTKO (black) and Ca_V_2 control (grey); Ca_V_2 control, 195 synapses/3 independent cultures; Ca_V_2 cTKO, 202/3; Ca_V_2 cTKO + Ca_V_2.1, 205/3; Ca_V_2 cTKO + Ca_V_1.3, 201/3. (F-H) As in C to E, but for synapses stained with antibodies against HA (to detect lentivirally expressed Ca_V_s, STED), PSD-95 (STED), and synapsin (confocal). Dashed lines in G denote levels in Ca_V_2 cTKO (black) and Ca_V_2 cTKO + Ca_V_2.1 (orange); Ca_V_2 control, 208/3; Ca_V_2 cTKO, 222/3; Ca_V_2 cTKO + Ca_V_2.1, 227/3; Ca_V_2 cTKO + Ca_V_1.3, 214/3. (I+J) Representative traces (I) and quantification (J) of NMDAR-mediated EPSCs recorded in whole-cell configuration and evoked by focal electrical stimulation; 18 cells/3 independent cultures each. (K+L) As in I and J, but for IPSCs; 18/3 each. Data are mean ± SEM; ***p < 0.001. Statistical significance compared to Ca_V_2 cTKO was determined with Kruskal-Wallis tests followed by Dunn’s multiple comparisons post-hoc tests for the proteins of interest or amplitudes in E, H, J and L. In H, the small but significant decreases in HA intensity in Ca_V_2 control and Ca_V_2 cTKO + Ca_V_1.3 compared to Ca_V_2 cTKO (which does not express an HA-tagged protein) are unlikely biologically meaningful. For C-terminal Ca_V_ sequences, and Ca_V_ expression analyses by Western blot and confocal microscopy, see Fig. S1.

For morphological analyses, neurons were stained with antibodies against Ca_V_2.1 or HA to detect Ca_V_s, PSD-95 to mark postsynaptic densities, and synapsin to label synaptic vesicle clusters. For STED analyses (Fig. 1C-H), we selected synapses in side-view through the presence of a vesicle cloud (imaged with confocal microscopy) and an elongated PSD-95 structure (STED) at one edge of the vesicle cloud, as established previously ^20,25,38,39,52^. We assessed Ca_V_ distribution and levels (STED) in these side-view synapses using line profiles drawn perpendicular to the PSD-95 structure, and we plotted the average line profiles (Fig. 1D+G) and peak intensities (Fig. 1E+H).

Endogenous and re-expressed Ca_V_2.1 formed elongated structures apposed to PSD-95 with a maximal intensity within tens of nanometers of the PSD-95 peak (Fig. 1C-H). We have established before that this distribution is characteristic of active zone localization ^20,25,39,53^. Furthermore, a strong PSD-95 peak was present in all conditions, matching our previous work that did not find morphological defects following Ca_V_2 triple knockout ^20^. Exogenously expressed Ca_V_1.3, monitored via the HA-tag, was not detected at the active zone (Fig. 1F-H). Consistent with the STED analyses, robust levels of Ca_V_2.1, but not Ca_V_1.3, were present in synaptic regions of interest (ROIs) defined by synapsin (Fig. S1F-I). Independent of their synaptic targeting, both Ca_V_2.1 and Ca_V_1.3 were effectively expressed in the somata of transduced Ca_V_2 cTKO neurons and in transfected HEK293T cells (Fig. S1C-E).

These morphological experiments were complemented with analyses of synaptic transmission in the same conditions (Fig. 1I-L). A focal stimulation electrode was used to evoke action potentials, and inhibitory or excitatory postsynaptic currents (IPSCs or EPSCs) were isolated pharmacologically. EPSCs were monitored via NMDA receptors because network excitation confounds the interpretation of EPSC amplitudes when AMPA receptors are not blocked. Ca_V_2 cTKO nearly abolished synaptic transmission, as characterized in detail before ^20^. Re-expression of Ca_V_2.1 restored EPSCs and IPSCs effectively, but exogenous expression of Ca_V_1.3 failed to produce any recovery (Fig. 1I-L), in agreement with the absence of Ca_V_1.3 from presynaptic sites (Fig. 1F-H). Taken together, these results establish that Ca_V_2.1, but not Ca_V_1.3, localizes to the active zone and gates neurotransmitter release when expressed in Ca_V_2 cTKO neurons.

### Ca_V_1.3 chimeras that contain the Ca_V_2.1 C-terminus localize to the active zone

Given the diverse interactions that converge within the Ca_V_ C-termini (Fig. S1A+B) ^20,42,43^, we hypothesized that the C-terminal sequences contain sufficient information to instruct Ca_V_ compartment specificity. To test this hypothesis, we generated two chimeric Ca_V_s: (1) in Ca_V_1.3, we replaced the entire intracellular C-terminus immediately after the last transmembrane segment with that of Ca_V_2.1, generating a channel we named Ca_V_1.3^2.1Ct^; and (2) we produced the inverse construct by replacing the Ca_V_2.1 C-terminus with that of Ca_V_1.3, generating Ca_V_2.1^1.3Ct^ (Figs. 2A, S1A). Both chimeric channels were efficiently expressed in transfected HEK293T cells (Fig. S2A) and were robustly detected in neuronal somata following lentiviral transduction of Ca_V_2 cTKO neurons (Fig. S2B+C).

**Figure 2.**
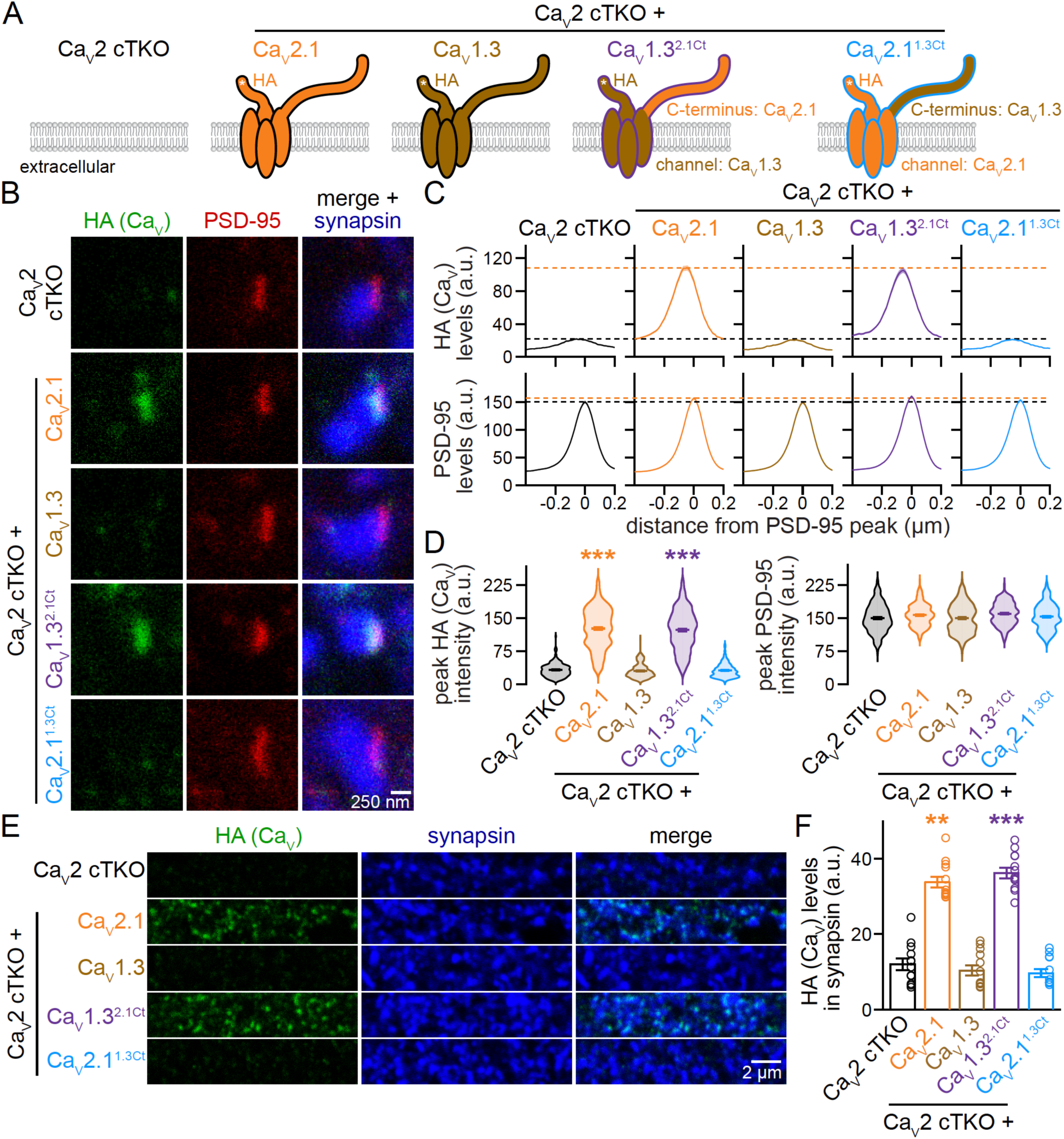
The Ca_V_2.1 C-terminus suffices to target Ca_V_1.3 to the presynaptic active zone. (A) Schematic of the conditions for comparison. (B-D) Representative images (B) and summary plots of intensity profiles (C) and peak levels (D) of HA and PSD-95 at side-view synapses stained for HA (STED), PSD-95 (STED), and synapsin (confocal). Dashed lines in C denote levels in Ca_V_2 cTKO (black) and Ca_V_2 cTKO + Ca_V_2.1 (orange); Ca_V_2 cTKO, 205 synapses/3 independent cultures; Ca_V_2 cTKO + Ca_V_2.1, 203/3; Ca_V_2 cTKO + Ca_V_1.3, 222/3; Ca_V_2 cTKO + Ca_V_1.3^2.1Ct^, 218/3; Ca_V_2 cTKO + Ca_V_2.1^1.3Ct^, 208/3. (E+F) Representative areas of confocal images (E) and quantification (F) of HA levels in synapsin regions of interest (ROIs). Identical areas (58.14 x 58.14 μm^2^) from the same cultures were imaged for confocal (E+F) and STED (B-D) analyses and whole images were quantified; 12 images/3 independent cultures each. Data are mean ± SEM; **p < 0.01 and ***p < 0.001. Statistical significance compared to Ca_V_2 cTKO was determined with Kruskal-Wallis tests followed by Dunn’s multiple comparisons post-hoc tests for the protein of interest in D and F. For Ca_V_ expression analyses by Western blot and confocal microscopy, see Fig. S2.

We then assessed the localization of these chimeric channels in the experimental setup described above and compared them side-by-side with Ca_V_2.1 and Ca_V_1.3. Strikingly, translocating the Ca_V_2.1 C-terminus onto Ca_V_1.3 efficiently targeted the resulting chimeric Ca_V_1.3^2.1Ct^ channel to the active zone in Ca_V_2 cTKO neurons, as assessed with STED microscopy (Fig. 2B-D). The distribution profile of Ca_V_1.3^2.1Ct^ and its abundance at the active zone recapitulated those of re-expressed Ca_V_2.1 (Fig. 2B-D). In contrast, the inverse swap abolished active zone localization of Ca_V_2.1^1.3Ct^ (Fig. 2B-D) despite effective somatic expression (Fig. S2B+C). Confocal microscopic analyses of Ca_V_ levels in synaptic ROIs corroborated these findings by revealing robust synaptic localization of Ca_V_1.3^2.1Ct^ but not of Ca_V_2.1^1.3Ct^ (Fig. 2E+F).

These results establish that Ca_V_1.3 is targeted to the presynaptic active zone when its C-terminus is replaced with that of Ca_V_2.1. Conversely, Ca_V_2.1 loses its active zone localization following the reverse swap. We conclude that the Ca_V_ C-termini contain sufficient information to define Ca_V_ compartment specificity, and these and previous data lead to two predictions. First, because removing known scaffolding motifs in the distal C-terminus only partially impaired active zone localization ^20,45^, there must be essential targeting motifs in the Ca_V_ C-terminus that have not yet been identified. Second, if the chimeric Ca_V_1.3^2.1Ct^ channel is appropriately coupled to primed vesicles within the active zone, then Ca_V_1.3^2.1Ct^ expression should restore synaptic transmission in Ca_V_2 cTKO neurons and render neurotransmitter release sensitive to L-type channel blockade. We next tested both predictions.

### An EF hand in the proximal C-terminus is necessary for Ca_V_2 active zone targeting

Removal of the known active zone scaffolding motifs in the Ca_V_2.1 C-terminus produces a partial defect in Ca_V_2.1 active zone targeting, but truncation of the entire C-terminus fully abolishes active zone localization ^20^. To define C-terminal sequences that contain unidentified targeting motifs, we segregated the Ca_V_2.1 C-terminus into a distal segment containing the active zone scaffolding motifs, and the complementary proximal segment (Fig. S1A+B). We generated two additional Ca_V_1.3 chimeras (Fig. 3A) by translocating either only the Ca_V_2.1 proximal C-terminus (Ca_V_1.3^2.1ProxCt^) or only the Ca_V_2.1 distal C-terminus (Ca_V_1.3^2.1DistCt^) onto Ca_V_1.3. Both Ca_V_1.3^2.1ProxCt^ and Ca_V_1.3^2.1DistCt^ were expressed efficiently in HEK293T cells after transfection (Fig. S3A) and in neuronal somata after lentiviral transduction (Fig. S3B+C). With STED microscopy, we detected Ca_V_1.3^2.1ProxCt^ at the active zone (Fig. 3B-D) of Ca_V_2 cTKO neurons. Active zone levels of Ca_V_1.3^2.1ProxCt^ were reduced compared to Ca_V_1.3^2.1Ct^ and resembled those of a mutant Ca_V_2.1 that lacks the active zone scaffolding motifs in the distal C-terminus ^20^. Hence, active zone targeting of chimeric Ca_V_1.3s operates in part through these distal sequences. Accordingly, Ca_V_1.3^2.1DistCt^ exhibited strong active zone localization in Ca_V_2 cTKO neurons and was indistinguishable from Ca_V_1.3^2.1Ct^ (Fig. 3B-D). Confocal analyses of protein levels in synaptic ROIs matched these findings (Fig. S3D+E).

**Figure 3.**
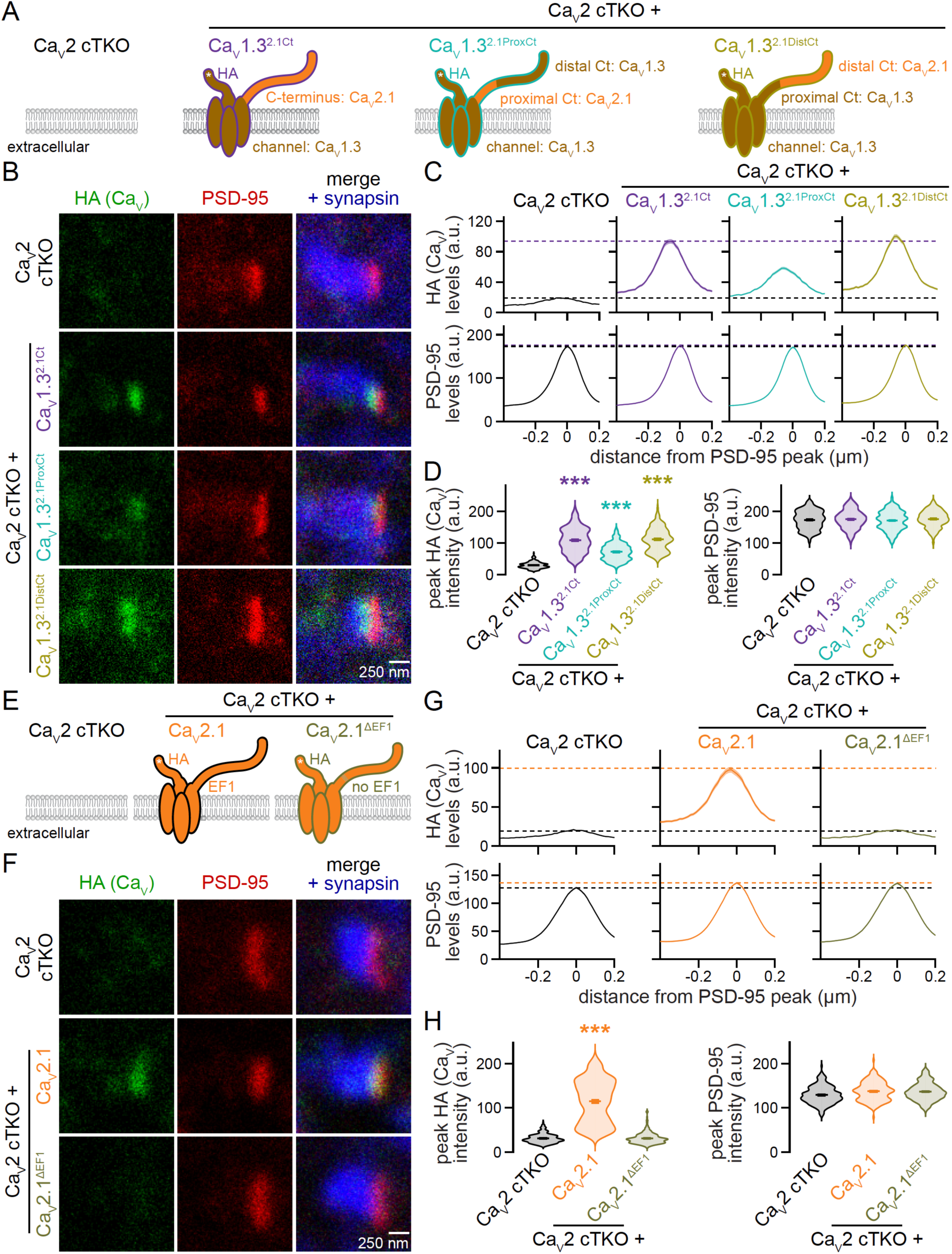
An EF hand in the proximal C-terminus is essential for Ca_V_2 active zone targeting. (A) Schematic of the conditions for comparison in B-D. (B-D) Representative images (B) and summary plots of intensity profiles (C) and peak levels (D) of HA and PSD-95 at side-view synapses stained for HA (STED), PSD-95 (STED), and synapsin (confocal). Dashed lines in C denote levels in Ca_V_2 cTKO (black) and Ca_V_2 cTKO + Ca_V_1.3^2.1Ct^ (purple); Ca_V_2 cTKO, 207 synapses/3 independent cultures; Ca_V_2 cTKO + Ca_V_1.3^2.1Ct^, 204/3; Ca_V_2 cTKO + Ca_V_1.3^2.1ProxCt^, 209/3; Ca_V_2 cTKO + Ca_V_1.3^2.1DistCt^, 210/3. (E) Schematic of the conditions for comparison in F-H. (F-H) Representative images (F) and summary plots of intensity profiles (G) and peak levels (H) of HA and PSD-95 at side-view synapses stained for HA (STED), PSD-95 (STED), and synapsin (confocal). Dashed lines in G denote levels in Ca_V_2 cTKO (black) and Ca_V_2 cTKO + Ca_V_2.1 (orange); Ca_V_2 cTKO, 200/3; Ca_V_2 cTKO + Ca_V_2.1, 180/3; Ca_V_2 cTKO + Ca_V_2.1^ΔEF1^, 203/3. Data are mean ± SEM; ***p < 0.001. Statistical significance compared to Ca_V_2 cTKO was determined with Kruskal-Wallis tests followed by Dunn’s multiple comparisons post-hoc tests for the protein of interest in D and H. For Ca_V_ expression analyses by Western blot and confocal microscopy, see Fig. S3.

Ca_V_1.3^2.1ProxCt^ demonstrates that translocation of the Ca_V_2.1 proximal C-terminus onto Ca_V_1.3 suffices to mediate some active zone localization (Fig. 3B-D) and indicates that the proximal C-terminal sequences are important for presynaptic trafficking. The Ca_V_ proximal C-termini (Fig. S1A+B) contain two EF hands ^54,55^. The first EF hand has been implicated in calmodulin-dependent modulation of Ca_V_ function ^56–58^, though no evidence to date establishes a role in Ca_V_ trafficking. We tested whether the first EF hand mediates active zone targeting by deleting the first EF hand from Ca_V_2.1 (Ca_V_2.1^ΔEF1^, Fig. 3E). Ca_V_2.1^ΔEF1^ was readily expressed in transfected HEK293T cells and detected in somata of lentivirally transduced neurons (Fig. S3F-H). However, deleting the first EF hand abolished Ca_V_2.1 active zone localization in STED microscopy (Fig. 3F-H) and rendered Ca_V_2.1^ΔEF1^ undetectable at synapses in confocal microscopy (Fig. S3I+J).

In summary, the Ca_V_2.1 distal C-terminus needs to be paired with proximal C-terminal elements to effectively localize Ca_V_s to the active zone. Our data establish that the proximal EF hand is required for active zone targeting of Ca_V_2.1.

### Ca_V_1.3^2.1Ct^ supports neurotransmitter release and confers L-type blocker sensitivity after Ca_V_2 ablation

Efficient neurotransmitter release requires that Ca_V_s are coupled to fusion-competent synaptic vesicles. Having demonstrated that translocation of the Ca_V_2.1 C-terminus directs Ca_V_1.3 to the active zone, we next asked whether the chimeric Ca_V_1.3^2.1Ct^ channel provides Ca^2+^ for action potential-triggered release (Fig. 4A). Ca_V_1.3^2.1Ct^ expression in Ca_V_2 cTKO neurons indeed resulted in EPSCs (Fig. 4B+C) and IPSCs (Fig. 4D+E) that were indistinguishable from those measured from Ca_V_2 cTKO neurons with re-expressed Ca_V_2.1. In contrast, and consistent with the loss of active zone targeting (Fig. 2B-D), Ca_V_2.1^1.3Ct^ failed to restore synaptic transmission (Fig. 4B-E).

**Figure 4.**
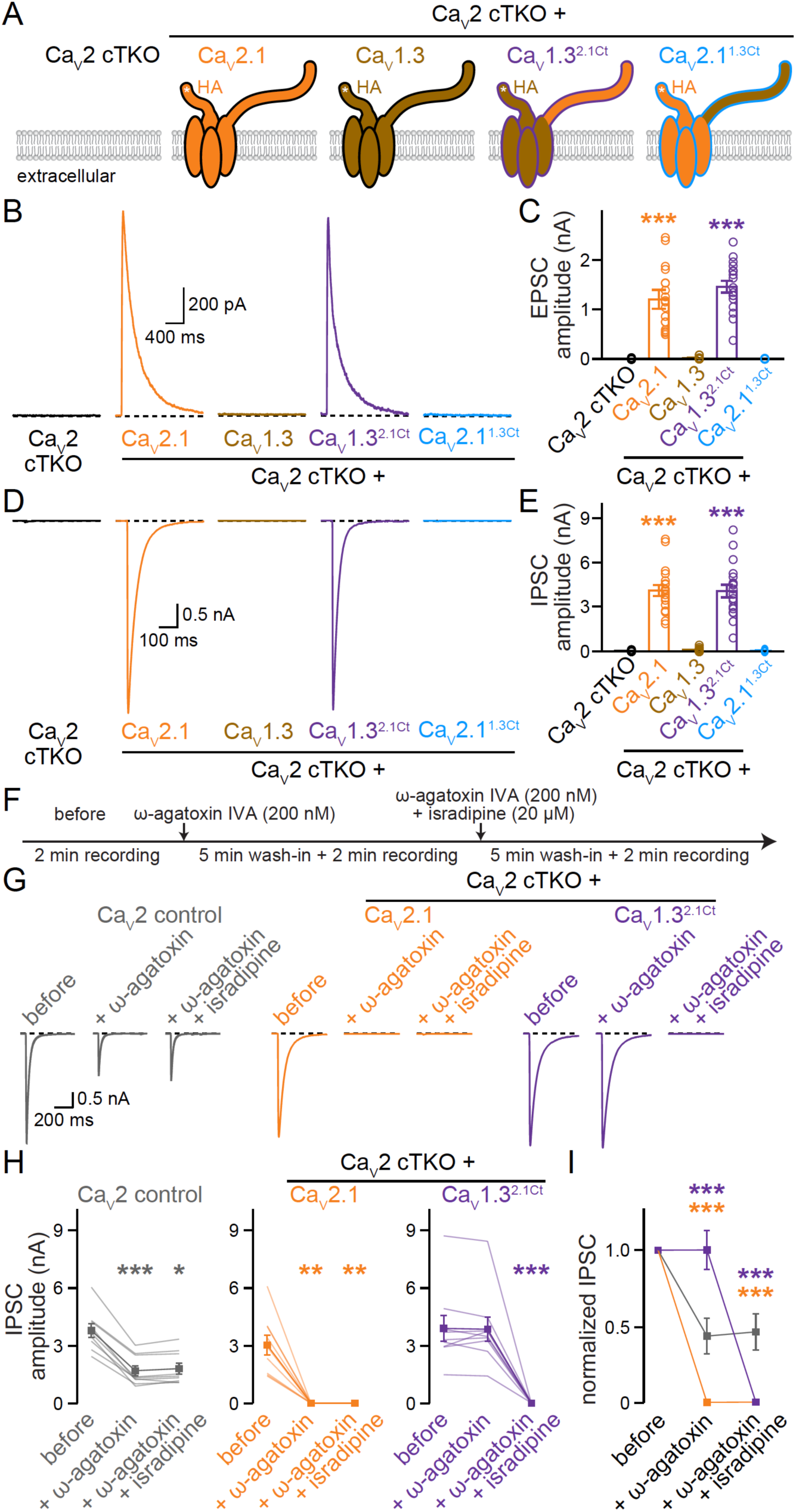
Ca_V_1.3^2.1Ct^ channels mediate neurotransmitter release and render it L-type blocker sensitive. (A) Schematic of the conditions for comparison, as in Fig. 2. (B+C) Representative traces (B) and quantification (C) of NMDAR-mediated EPSCs; 18 cells/3 independent cultures each. (C+E) As in B and C, but for IPSCs; 18/3 each. (F) Experimental strategy to evaluate blocker sensitivity of synaptic transmission. Evoked IPSCs were recorded before blocker application (before), after wash-in of 200 nM ω-agatoxin-IVA alone (+ ω-agatoxin, to block Ca_V_2.1), and after wash-in 200 nM ω-agatoxin-IVA and 20 μM isradipine (+ ω-agatoxin + isradipine, to block Ca_V_1s and Ca_V_2.1). (G+H) Representative traces (G) and quantification (H) of IPSCs recorded as outlined in F; 9 cells/3 independent cultures each. (I) Comparison of IPSCs normalized to “before” in each condition; 9/3 each. Data are mean ± SEM; *p < 0.05, **p < 0.01, and ***p < 0.001. Statistical significance compared to Ca_V_2 cTKO in C and E was determined with Kruskal-Wallis tests followed by Dunn’s multiple comparisons post-hoc tests. Statistical significance compared to “before” in H was determined with Friedman tests followed by Dunn’s multiple comparisons post-hoc tests. Statistical significance compared to Ca_V_2 control in I was determined with two-way, repeated-measures ANOVA followed by Dunnett’s multiple comparisons post-hoc tests. For characterization of C-terminally truncated Ca_V_1.3, see Fig. S4; for assessment of isradipine-sensitivity of synaptic transmission in Ca_V_2 control neurons, see Fig. S5.

It is possible that the presynaptic targeting and function of Ca_V_1.3^2.1Ct^ results from removal of a dendritic targeting sequence rather than addition of an axonal targeting motif. To address this possibility, we generated a Ca_V_1.3 lacking the entire C-terminus (Ca_V_1.3^ΔCt^). Ca_V_1.3^ΔCt^ was effectively expressed (Fig. S4A-D) but was not targeted to synapses (Fig. S4E+F) or active zones (Fig. S4G-I). Furthermore, Ca_V_1.3^ΔCt^ did not mediate neurotransmitter release (Fig. S4J-M). We conclude that active zone targeting of Ca_V_1.3^2.1Ct^ arises from an instructive role of the Ca_V_2.1 C-terminus.

At central synapses, neurotransmitter release is insensitive to L-type Ca_V_ blockade (Fig. S5) ^17^. Given that we replaced presynaptic Ca_V_2s with an L-type-like Ca_V_ (Ca_V_1.3^2.1Ct^), we finally tested whether we also altered the pharmacological sensitivity of synaptic transmission. We performed serial Ca_V_ blockade (Fig. 4F) through sequential application of ω-agatoxin IVA (to block Ca_V_2.1) and isradipine (to block Ca_V_1s). In Ca_V_2 control neurons, ω-agatoxin IVA reduced IPSCs approximately by half (Fig. 4G-I), consistent with the reliance of neurotransmitter release on both Ca_V_2.1 and Ca_V_2.2 ^24,59^. Isradipine had no effect in Ca_V_2 control neurons (Fig. S5). Naturally, ω-agatoxin IVA fully inhibited synaptic transmission in Ca_V_2 cTKO neurons rescued with Ca_V_2.1. However, for Ca_V_2 cTKO neurons that expressed Ca_V_1.3^2.1Ct^, synaptic transmission was resistant to ω-agatoxin IVA and instead wholly sensitive to the L-type channel blocker isradipine (Fig. 4G-I). Hence, Ca_V_1.3^2.1Ct^ functionally replaces endogenous Ca_V_2s in Ca_V_2 cTKO neurons and renders neurotransmission fully dependent on L-type Ca_V_ activity.

## Discussion

Voltage-gated Ca^2+^ channels are a prototypical protein family to illustrate neuronal polarization: distinct Ca_V_s are sorted effectively to dendritic, somatic and axonal compartments. Here, we establish that the Ca_V_ C-termini contain the necessary and sufficient information to sort Ca_V_s into specific subcellular compartments. Within the C-terminus of Ca_V_2.1, the proximal EF hand is essential for presynaptic targeting and it operates in concert with distal scaffolding motifs. Together, the Ca_V_2.1 C-terminal sequences are sufficient to re-direct somatodendritic Ca_V_1 channels to the active zone. Conversely, the Ca_V_1.3 C-terminal sequences disrupt Ca_V_2.1 active zone localization. Our work establishes mechanisms for compartment-specific targeting of a protein family central to the polarized organization of neurons.

Multiple cargo selectivity filters converge within the endoplasmic reticulum, the Golgi apparatus, the axon initial segment, and the presynaptic bouton that together permit the targeting of a limited subset of proteins to the active zone while deflecting other cargo ^60,61^. Sequence motifs within these proteins may dictate compartment sorting at two major checkpoints: (1) they may mediate protein recruitment into cargo vesicles that are directed to the axon, and (2) they may stabilize proteins at the active zone following their delivery ^2,62^. Our work establishes that the Ca_V_2.1 C-terminus encodes necessary and sufficient information to navigate these two checkpoints and implies a cooperative relationship between the proximal and distal elements. The Ca_V_2.1 distal C-terminus efficiently localizes chimeric Ca_V_1.3s to the active zone, indicating that the distal C-terminal sequences permit both Ca_V_ sorting into presynaptic cargo and Ca_V_ tethering at the active zone, so long as a proximal EF hand is present. The distal motifs that bind to active zone proteins likely fulfill these roles as disrupting their interactions with RIM and RIM-BP leads to targeting defects ^20,24,36,37,45^ similar to those exhibited by chimeric Ca_V_1.3s with the Ca_V_2.1 proximal C-terminus and the Ca_V_1.3 distal C-terminus (Fig. 3).

The efficiency with which the chimeric Ca_V_1.3^2.1Ct^ and Ca_V_1.3^2.1DistCt^ channels are targeted to the active zone establishes that the proximal C-termini of both Ca_V_1.3 and Ca_V_2.1 contain necessary information for active zone Ca_V_ delivery. This is in line with the high homology of the EF hands and IQ-motif across Ca_V_ proximal C-termini and with the presence of these sequences in other voltage-gated channels ^55,63^. The proximal C-terminus might include multiple instructive signals that together inform Ca_V_ targeting. The EF hand binds to AP-1 and possibly Ca^2+^, which could provide for a trafficking control checkpoint ^64,65^. Calmodulin binds to the IQ motif and might regulate channel trafficking and function ^10,33,34,56^. Other unknown interactions with these sequences or with sequences elsewhere in the proximal C-terminus might be involved in targeting as well. Altogether, we posit that the proximal EF hand is necessary for passing a trafficking checkpoint that permits incorporation of these Ca_V_s into axon-bound cargo, but likely has no role in stabilizing Ca_V_s within the active zone.

Our work on Ca_V_s provides mechanistic insight into the polarized trafficking of protein material in neurons and raises multiple questions. First, some synapses depend on only a single Ca_V_2 subtype while others redundantly use multiple Ca_V_2s, and some synapses experience developmental switches in their Ca_V_2 usage ^66,67^. Whether there are specific trafficking and anchoring mechanisms or whether these properties are determined wholly by switches in gene expression remains to be determined. Second, the proximal sequences we identified as important for targeting are also present in other ion channels that undergo polarized trafficking, for example in neuronal Na^+^ channels ^55,63^. It is possible that the mechanisms we describe for Ca_V_s are broadly employed across channel proteins. The example of Ca_V_s forms an ideal framework to build on and further define mechanisms that sort proteins into specific neuronal compartments.

## Acknowledgements

We thank members of the Kaeser laboratory for insightful discussions and comments, and we specifically acknowledge R. Held, H. Nyitrai, C. Tan, K. Ma, and A. Morabito for help and advice early in the project and/or feedback on the findings and the manuscript. We acknowledge J. Wang, C. Qiao, V. Charles and G. Handy for technical support. We thank A.M.J.M. van den Maagdenberg for providing Ca_V_2.1 floxed mice, and T. Schneider for Ca_V_2.3 floxed mice. This work was supported by the NIH (R01NS083898 to PSK, F31NS127399 to MC), by a Stuart H.Q. and Victoria Quan fellowship (to MC), and by Harvard Medical School. We acknowledge the Neurobiology Imaging Facility at Harvard Medical School for microscope availability and support.

## Author contributions

Conceptualization, MC and PSK; Methodology, MC; Formal Analysis, MC and PK; Investigation, MC; Resources, MC; Writing-Original Draft, MC and PSK; Writing-Review & Editing, MC and PSK; Supervision, PSK; Funding Acquisition, PSK.

---

**Figure S1.**
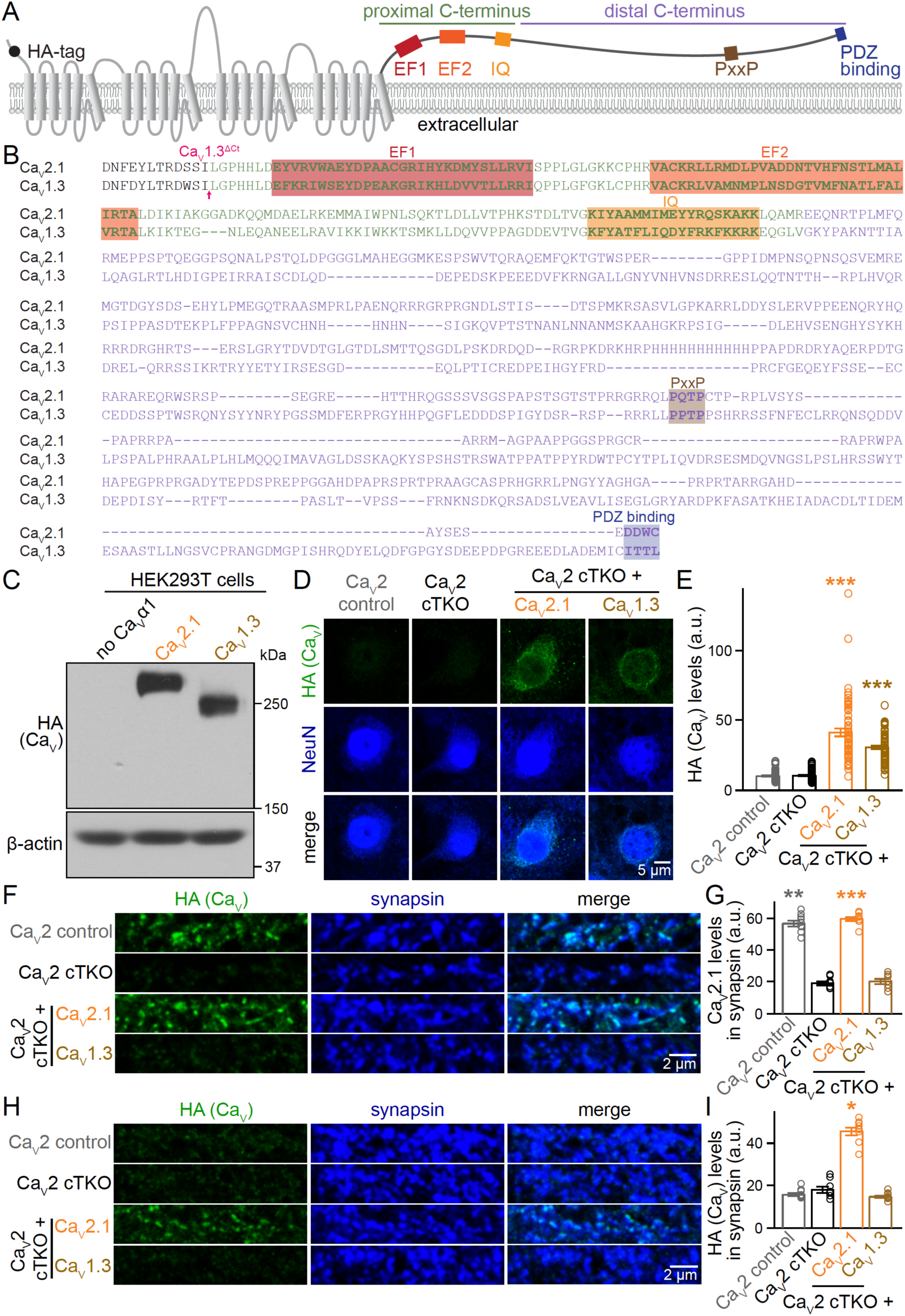
Additional assessment of Ca_V_2.1 and Ca_V_1.3 expression and localization. (A) Schematic of Ca_V_2.1 with important sequence motifs highlighted, adapted from ^20^; EF1 and EF2: EF hands; IQ: IQ motif; PxxP: proline rich motif. (B) Alignment of the C-terminal sequences starting immediately after the last transmembrane segment (for Ca_V_2.1, residues DNFE…DDWC are matching with GenBank Entry AY714490.1; for Ca_V_1.3, residues DNFD…ITTL are matching with GenBank Entry AF370010.1). Sequence motifs are highlighted, and the Ca_V_ proximal and distal C-terminal segments are labeled in green and purple, respectively. (C) Western blot of HEK293T cell homogenates after transfection with Ca_V_β1, Ca_V_α2δ1, and without (no Ca_V_α1) or with a Ca_V_α1 subunit to assess Ca_V_α1 expression; Ca_V_2.1 and Ca_V_1.3 were transfected and analyzed multiple times, but only once in this order. (D+E) Representative confocal images (D) and quantification (E) of HA levels in cell bodies of neurons stained with antibodies against HA and NeuN. Cell bodies were defined as donut shaped ROIs using the outer edge of the NeuN profile along the main somatic compartment not including the neurites, and by excluding the EGFP-labeled nucleus; 60 somata/3 independent cultures each. (F+G) Representative areas of confocal images (F) and quantification (G) of Ca_V_2.1 levels in synapsin ROIs (the imaged areas are identical to the STED scans in Fig. 1C-E); Ca_V_2 control, 9 images/3 independent cultures; Ca_V_2 cTKO, 8/3; Ca_V_2 cTKO + Ca_V_2.1, 9/3; Ca_V_2 cTKO + Ca_V_1.3, 8/3. (H+I) As in F and G, but for neurons stained with antibodies against HA, PSD-95 and synapsin (the imaged areas are identical to the STED scans in Fig. 1F-H); Ca_V_2 control, 9/3; Ca_V_2 cTKO, 8/3; Ca_V_2 cTKO + Ca_V_2.1, 9/3; Ca_V_2 cTKO + Ca_V_1.3, 9/3. Data are mean ± SEM; *p < 0.05, **p < 0.01, and ***p < 0.001. Statistical significance compared to Ca_V_2 cTKO was determined with Kruskal-Wallis tests followed by Dunn’s multiple comparisons post-hoc tests for the proteins of interest in E, G, and I.

**Figure S2.**
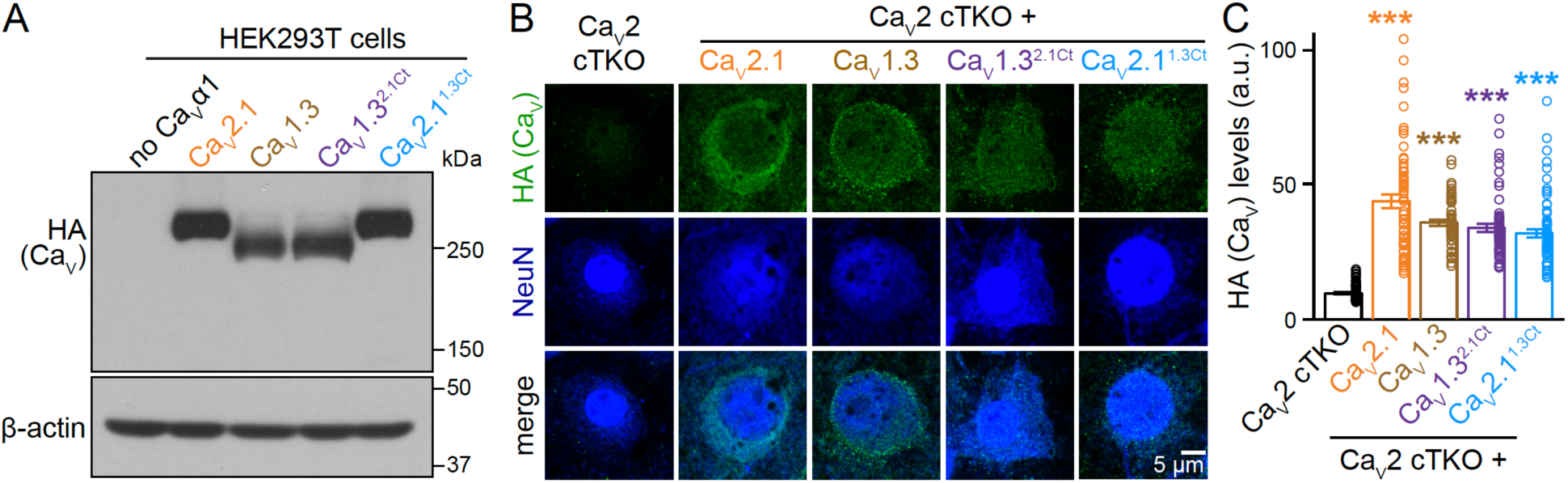
Additional assessment of Ca_V_1.3^2.1Ct^ and Ca_V_2.1^1.3Ct^ expression. (A) Western blot of HEK293T cell homogenates after transfection with Ca_V_β1, Ca_V_α2δ1, and without (no Ca_V_α1) or with a Ca_V_α1 subunit to assess Ca_V_α1 expression, a representative blot from three independent repeats is shown. (B+C) Representative confocal images (B) and quantification (C) of HA levels in cell bodies of neurons stained with antibodies against HA and NeuN; 60 somata/3 independent cultures each. Data are mean ± SEM; ***p < 0.001. Statistical significance compared to Ca_V_2 cTKO was determined with Kruskal-Wallis tests followed by Dunn’s multiple comparisons post-hoc tests for the protein of interest in C.

**Figure S3.**
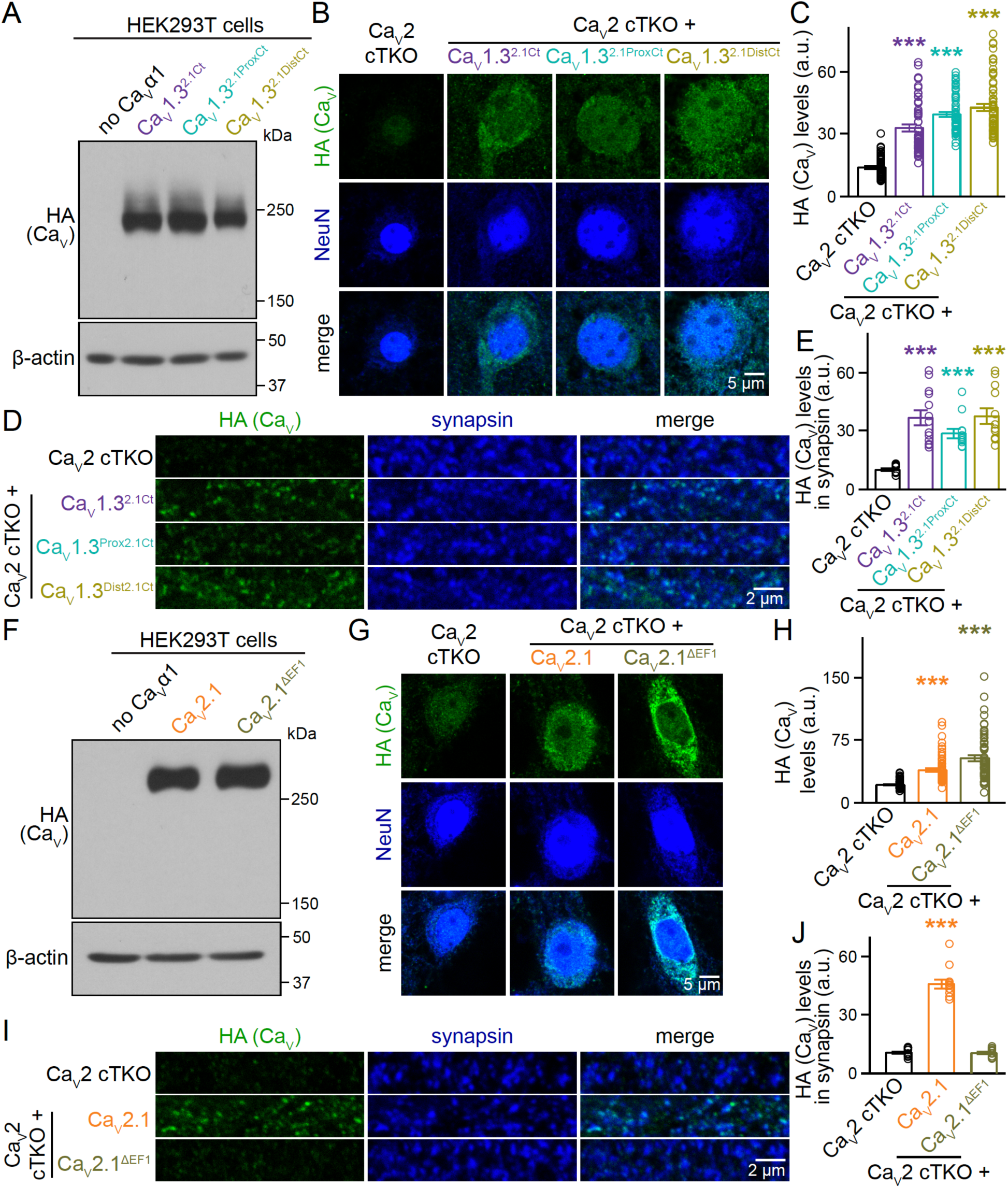
Additional assessment of Ca_V_1.3^2.1ProxCt^, Ca_V_1.3^2.1DistCt^ and Ca_V_2.1^ΔEF1^ expression and localization. (A) Western blot of HEK293T cell homogenates after transfection with Ca_V_β1, Ca_V_α2δ1, and without (no Ca_V_α1) or with a Ca_V_α1 subunit to assess Ca_V_α1 expression, a representative blot from three independent repeats is shown. (B+C) Representative confocal images (B) and quantification (C) of HA levels in cell bodies of neurons stained with antibodies against HA and NeuN; 60 somata/3 independent cultures each. (D and E) Representative areas of confocal images (D) and quantification (E) of HA levels in synapsin ROIs (the imaged areas are identical to the STED scans in Fig. 3B-D); Ca_V_2 control, 9 images/3 independent cultures; Ca_V_2 cTKO, 12/3; Ca_V_2 cTKO + Ca_V_1.3^2.1Ct^, 13/3; Ca_V_2 cTKO + Ca_V_1.3^2.1ProxCt^, 12/3; Ca_V_2 cTKO + Ca_V_1.3^2.1DistCt^, 12/3. (F) Western blot of HEK293T cell homogenates after transfection with Ca_V_β1, Ca_V_α2δ1, and without (no Ca_V_α1) or with a Ca_V_α1 subunit to assess expression, a representative blot from two independent repeats is shown. (G+H) Representative confocal images (G) and quantification (H) of HA levels in cell bodies of neurons stained with antibodies against HA and NeuN; 60/3 each. (I and J) Representative areas of confocal images (I) and quantification (J) of HA levels in synapsin ROIs (the imaged areas are identical to the STED scans in Fig. 3F-H); 12/3 each. Data are mean ± SEM; ***p < 0.001. Statistical significance compared to Ca_V_2 cTKO was determined with Kruskal-Wallis tests followed by Dunn’s multiple comparisons post-hoc tests for the protein of interest in C, E, H, and J.

**Figure S4.**
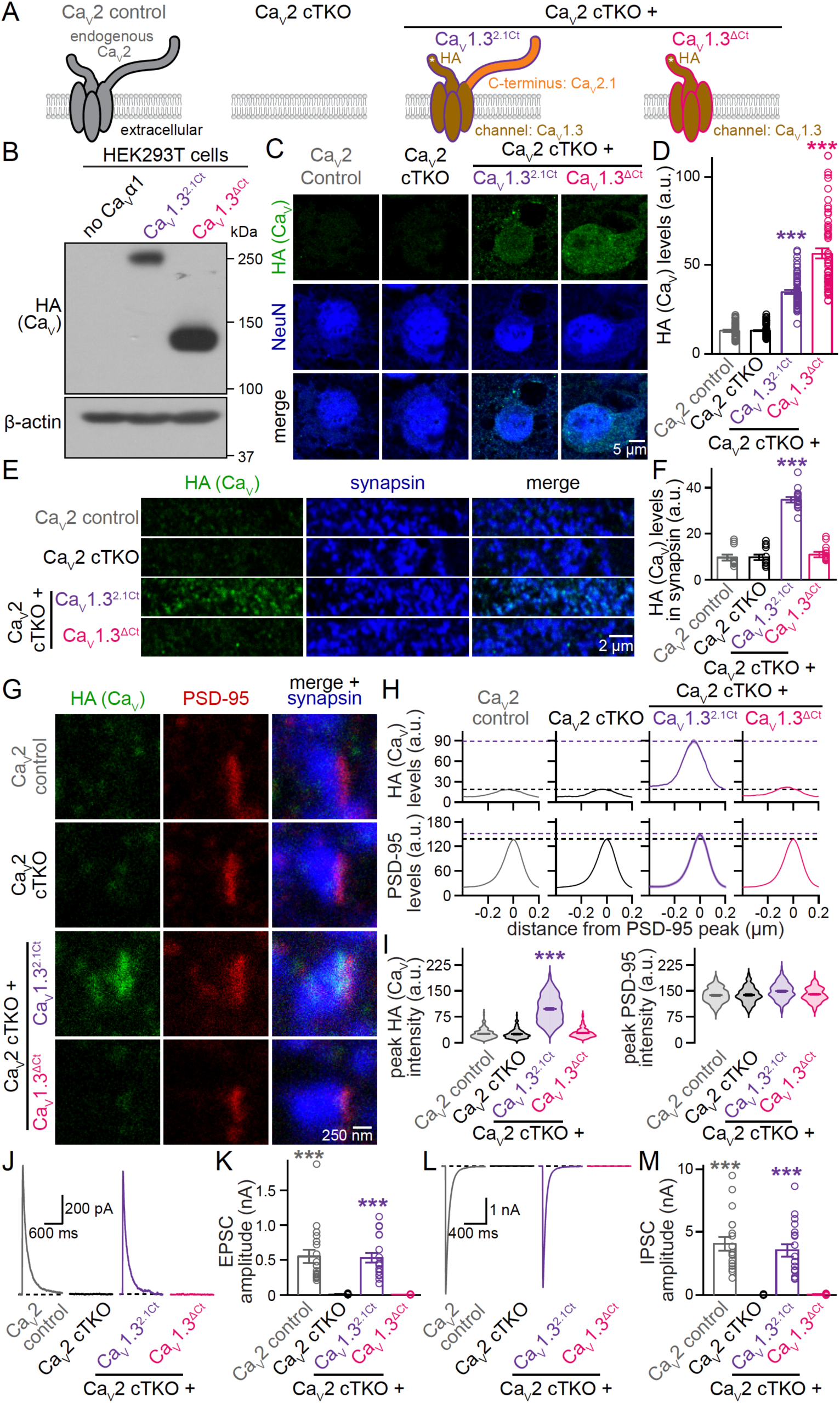
Assessment of Ca_V_1.3^ΔCt^. (A) Schematic of the conditions for comparison. (B) Western blot of HEK293T cell homogenates after transfection with Ca_V_β1, Ca_V_α2δ1, and without (no Ca_V_α1) or with a Ca_V_α1 subunit to assess Ca_V_α1 expression, a representative blot from two independent repeats is shown. (C+D) Representative confocal images (C) and quantification (D) of HA levels in cell bodies of neurons stained with antibodies against HA and NeuN; 60 somata/3 independent cultures each. (E+F) Representative areas of confocal images (E) and quantification (F) of HA levels in synapsin ROIs; Ca_V_2 control, 11 images/3 independent cultures; Ca_V_2 cTKO, 12/3; Ca_V_2 cTKO + Ca_V_1.3^2.1Ct^, 14/3; Ca_V_2 cTKO + Ca_V_1.3^ΔCt^, 14/3. (G-I) Representative images (G) and summary plots of intensity profiles (H) and peak levels (I) of HA and PSD-95 at side-view synapses stained for HA (STED), PSD-95 (STED), and synapsin (confocal). The imaged areas are identical to the ones used for confocal analyses in E+F. Dashed lines in H denote levels in Ca_V_2 cTKO (black) and Ca_V_2 cTKO + Ca_V_1.3^2.1Ct^ (purple); Ca_V_2 control, 198 synapses/3 independent cultures; Ca_V_2 cTKO, 190/3; Ca_V_2 cTKO + Ca_V_1.3^2.1Ct^, 207/3; Ca_V_2 cTKO + Ca_V_1.3^ΔCt^, 195/3. (J+K) Representative traces (J) and quantification (K) of NMDAR-mediated EPSCs; 18 cells/3 independent cultures each. (L+M) As in J and K, but for IPSCs; 18/3 each. Data are mean ± SEM; and ***p < 0.001. Statistical significance compared to Ca_V_2 cTKO was determined with Kruskal-Wallis tests followed by Dunn’s multiple comparisons post-hoc tests for the protein of interest or amplitudes in D, F, I, K, and M.

**Figure S5.**
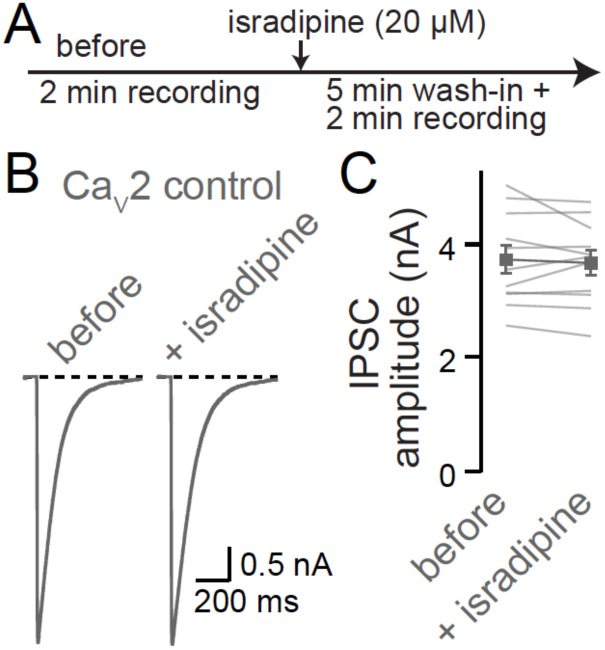
Assessment of L-type blocker sensitivity of synaptic transmission in Ca_V_2 control neurons. (A) Experimental strategy to evaluate blocker sensitivity of synaptic transmission. (B+C) Representative traces (B) and quantification (C) of IPSCs recorded as outlined in A; 11 cells/3 independent cultures. Data are mean ± SEM.

## Materials and methods

### Mice

Ca_V_2 conditional triple homozygote floxed mice were described before ^20^ and they contain homozygote floxed alleles for Ca_V_2.1 (*Cacna1a*, ^68^), Ca_V_2.2 (*Cacna1b*, ^20^), and Ca_V_2.3 (*Cacna1e*, ^69^). Mice were housed as breeding pairs or separated by sex, and they were under a 12 h light-dark cycle with free access to food and water in a room set to 22 °C (range 20-24 °C) and 50% humidity (range 35-70%). Mice were genotyped either in the lab following established protocols ^20^ or by Transnetyx. For *Cacna1a*, the following oligonucleotide primer pair was used for in-lab genotyping: forward, ACCTACAGTCTGCCAGGAG; reverse, TGAAGCCCAGACATCCTTGG (expected band sizes, wild type: 393 bp, floxed: 543 bp); for *Cacna1b*: forward, TGGTTGGTGTCCTGTTCTCC; reverse, TAAGGAGCAGGGAATCCTGG (expected band sizes, wild type: 219bp, floxed: 359 bp); for *Cacna1c*: forward, GACAAGACCCCAATGTCTCG; reverse, TCCATGTTCCTTCTCACTCC (expected band sizes, wild type: 295 bp, floxed: 334 bp). Animal experiments were performed according to approved protocols at Harvard University.

### Primary neuronal cultures

Primary mouse hippocampal cultures were generated from newborn mice as described previously ^20,38,39^. Hippocampi were dissected out from newborn mice within 24 h after birth. Cells were dissociated and plated onto Matrigel-treated glass coverslips in plating medium composed of Minimum Essential Medium (MEM) with 0.5% glucose, 0.02% NaHCO3, 0.1 mg/mL transferrin, 10% Fetal Select bovine serum (Atlas Biologicals FS-0500-AD), 2 mM L-glutamine, and 25 μg/mL insulin. Cells from mice of both sexes were mixed. Cultures were maintained in a 37 °C-tissue culture incubator, and after ∼24 h the plating medium was exchanged with growth medium composed of MEM with 0.5% glucose, 0.02% NaHCO3, 0.1 mg/mL transferrin, 5% Fetal Select bovine serum (Atlas Biologicals FS-0500-AD), 2% B-27 supplement (Thermo Fisher 17504044), and 0.5 mM L-glutamine. On day in vitro (DIV) 1 to 2, depending on growth, 50% or 75% of the medium was exchanged with growth medium supplemented with 4 μM Cytosine β-D-arabinofuranoside (AraC). Experiments and analyses were performed at DIV15 to 19, as described below.

### Cell lines

HEK293T cells, an immortalized cell line of female origin, were cultured as described before ^20,38,39^. They were purchased from ATCC (CRL-3216, RRID: CVCL_0063), expanded, and stored in liquid nitrogen until use. After thawing, the cells were grown in Dulbecco’s Modified Eagle Medium (DMEM) with 10% fetal bovine serum (Atlas Biologicals F-0500-D) and 1% penicillin-streptomycin. HEK293T cells were passaged every 1 to 3 d at a ratio of 1:3 to 1:10. HEK293T cell batches were typically replaced after 20 passages by thawing a fresh vial from the expanded stock.

### Lentiviruses

Lentiviruses used to transduce primary hippocampal neurons were produced in HEK293T cells. HEK293T cells were transfected with the Ca^2+^ phosphate method with REV (p023), RRE (p024) and VSVG (p025), as well as a lentiviral plasmid encoding the protein of interest. For Ca_V_ proteins of interest, these were plasmids p789, p947, p1077, p1078, p1079, p1080, p1083, and p1084. To produce lentiviruses expressing EGFP-tagged Cre recombinase (to generate Ca_V_2 cTKO neurons), pFSW EGFP-Cre (p009) was used. For lentiviruses expressing a truncated, enzymatically inactive EGFP-tagged Cre (to generate Ca_V_2 control neurons), pFSW EGFP-ΔCre (p010) was used. Plasmids were transfected at a 1:1:1:1 molar ratio and with a total amount of 6.7 μg DNA. Approximately 24 h after transfection, the medium was switched to neuronal growth medium (described above), and the HEK293T cell supernatant was harvested 24-36 h later by centrifugation at 700 x g. For expression of EGFP-Cre and EGFP-ΔCre, neurons were infected by adding HEK293T cell supernatant at DIV5. For expression of Ca_V_s, neurons were infected at DIV1. Ca_V_2 control neurons were additionally infected with a virus made using a pFSW plasmid (p008) lacking a cDNA in the multiple cloning site in place of an expression virus. Neuronal protein expression from these lentiviruses was driven by a human synapsin promoter ^38,70^.

### Ca_V_ expression constructs

For experiments in neurons, lentiviral backbones containing a human synapsin promoter were used (pFSW HA-Ca_V_2.1, p789; pFSW HA-Ca_V_1.3, p1077; pFSW HA-Ca_V_1.3^2.1Ct^, p1078; pFSW HA-Ca_V_2.1^1.3Ct^, p1079; pFSW HA-Ca_V_1.3^ΔCt^, p1080; pFSW HA-Ca_V_1.3^2.1ProxCt^, p1083; pFSW HA-Ca_V_1.3^2.1DistCt^, p1084; pFSW HA-Ca_V_2.1^ΔEF1^, p947). For experiments in HEK293T cells, expression vectors with a CMV promoter were used (pCMV HA-Ca_V_2.1, p771; pCMV HA-Ca_V_1.3, p1073; pCMV HA-Ca_V_1.3^2.1Ct^, p1074; pCMV HA-Ca_V_2.1^1.3Ct^, p1075; pCMV HA-Ca_V_1.3^ΔCt^, p1076; pCMV HA-Ca_V_1.3^2.1ProxCt^, p1081; pCMV HA-Ca_V_1.3^2.1DistCt^, p1082; pCMV HA-Ca_V_2.1^ΔEF1^, p939). For these constructs, the Ca_V_ coding sequences were identical between corresponding pFSW and pCMV versions. The <u>sequence of Ca_V_2.1</u> was identical to GenBank Entry AY714490.1 (mouse) with the addition of an HA-tag after position V_27_ flanked by short, exogenous linkers. The resulting cDNAs (p771 and p789) had the sequence M_1_ARF…GVVV_27_-AS-YPYDVPDYA-ACR-G_28_AAG…DDWC_2369_. The <u>sequence of Ca_V_1.3</u> was as follows: the pore region was identical to residues M_1_QHQ…FDYL_1466_ from Ca_V_1.3e[8a,11,31b,Δ32,42a] (rat) and corresponds to residues M_10_QHQ…FDYL_1475_ of GenBank Entry EDL89004.1.

Ca_V_1.3e[8a,11,31b,Δ32,42a] was a gift from D. Lipscombe (Addgene Plasmid #49333; http://n2t.net/addgene:49333; RRID:Addgene_49333) ^71^. The intracellular C-terminal tail was identical to residues T_7_ to L_695_ from GenBank Entry AF370010.1 (a partial cDNA, rat); a Ca_V_1.3 plasmid containing this C-terminal tail was a gift from I. Bezprozvanny ^43^. An HA-tag was inserted after position G_29_ (referring to the numbering of Addgene Plasmid #49333) and flanked by short, exogenous linkers. The resulting cDNAs (p1073 and p1077) had the sequence M_1_QHQ…SGEG_29_-AS-YPYDVPDYA-ACR-P_30_TSQ…FDYL_1466_-T_1467_RDW…ITTL_2155_, with M_1_QHQ-SGEG_29_ and P_30_TSQ-FDYL_1466_ derived from Addgene Plasmid #49333 ^71^, and with T_1467_RDW-ITTL_2155_ derived from the plasmid obtained from I. Bezprozvanny ^43^. The <u>sequence of Ca_V_1.3^2.1Ct^</u> (p1074 and p1078) contained the pore region (MQHQ…DWSI) from p1077 (Ca_V_1.3) followed by the C-terminus (LGPH…DDWC) from p789 (Ca_V_2.1, see Fig. S1B). The <u>sequence of Ca_V_2.1^1.3Ct^</u> (p1075 and p1079) contained the pore region (MARF…FEYL) from p789 (Ca_V_2.1) followed by the C-terminus (TRDW…ITTL) from p1077 (Ca_V_1.3, see Fig. S1B). The <u>sequence of Ca_V_1.3^2.1ProxCt^</u> (p1081 and 1083) contained the pore region (MQHQ…DWSI) from p1077 (Ca_V_1.3), followed by the proximal C-terminus (LGPH…QAMR) from p789 (Ca_V_2.1) and then by the distal C-terminus (GKYP…ITTL) from p1077 (Ca_V_1.3, see Fig. S1B). The <u>sequence of Ca_V_1.3^2.1DistCt^</u> (p1082 and 1084) contained the pore region and the proximal C-terminus (MQHQ…QGLV) from p1077 (Ca_V_1.3) followed by the distal C-terminus (EEQN…DDWC) from p789 (Ca_V_2.1, see Fig. S1B). In the sequence of Ca_V_2.1^ΔEF1^ (p939 and p947), the first EF hand (EYVR…LLRVI) was replaced with residues EY in p789 (Ca_V_2.1, see Fig. S1B). The <u>sequence of Ca_V_1.3^ΔCt^</u> (p1076 and 1080) contained the pore region (MQHQ…DWSI) from p1077 (Ca_V_1.3) and did not contain a C-terminus (see Fig. S1B).

### Confocal and STED microscopy of synapses

Confocal and STED microscopy and analyses were performed as described before ^20,25,39,53,72,73^. Neurons cultured on 0.17 mm thick 12 mm diameter (#1.5) coverslips were washed two times with PBS warmed to 37 °C, and then fixed in 2% PFA + 4% sucrose (in PBS) at room temperature. After fixation, coverslips were rinsed three times in PBS + 50 mM glycine, then permeabilized in PBS + 0.1% Triton X-100 + 3% BSA (TBP) for 1 h at room temperature. Coverslips were stained with primary antibodies diluted in TBP for ∼48 h at 4 °C. The following primary antibodies were used: mouse IgG1 anti-HA (1:500, RRID: AB_2565006, A12), rabbit anti-Ca_V_2.1 (1:200, RRID: AB_2619841, A46), guinea pig anti-PSD-95 (1:500, RRID: AB_2619800, A5), rabbit anti-synapsin (1:500, RRID: AB_2200097, A30), and mouse IgG1 anti-synapsin (1:500, RRID_2617071, A57). After primary antibody staining, coverslips were rinsed twice and washed three times for 5 min in PBS + 50 mM glycine at room temperature. Alexa Fluor 488 (to detect HA-tagged Ca_V_s or endogenous Ca_V_2.1; anti-mouse IgG1, RRID: AB_2535764, S7; or, anti-rabbit, RRID: AB_2576217, S5), Alexa Fluor 555 (to detect the postsynaptic marker PSD-95; anti-guinea pig, RRID: AB_2535856, S23), and Alexa Fluor 633 (to detect the synaptic vesicle cloud; anti-rabbit, RRID: AB_2535731, S33; or, anti-mouse IgG1, RRID: AB_2535768, S29) conjugated antibodies were diluted in TBP at 1:200 (for Alexa Fluor 488 and 555) or 1:500 (for Alexa Fluor 633), and coverslips were incubated with the secondary antibody solution for ∼24 h at 4 °C. Coverslips were then rinsed twice with PBS + 50 mM glycine and once with deionized water, air-dried and mounted on glass slides in fluorescent mounting medium. Confocal and STED images were acquired on a Leica SP8 Confocal/STED 3X microscope with an oil immersion 100x 1.44 numerical aperture objective and gated detectors as described previously ^20,72^. 58.14 x 58.14 μm^2^ areas were acquired using 2x digital zoom (4096 x 4096 pixels, pixel size of 14.194 x 14.194 nm^2^). Alexa Fluor 633, Alexa Fluor 555, and Alexa Fluor 488 were excited at 633 nm, 555 nm and 488 nm using a white light laser at 1-10% of 1.5 mW laser power. The Alexa Fluor 633, Alexa Fluor 555, and Alexa Fluor 488 channels were acquired first in confocal mode. For the Alexa Fluor 555 and Alexa Fluor 488 channels, the same areas were then sequentially acquired in STED mode using 660 nm and 592 nm depletion lasers, respectively. Identical imaging and laser settings were applied to all conditions within a given biological repeat. For analyses of presynaptic Ca_V_ distribution in STED images, synapses were selected in side-view. Side-view synapses were defined as synapses that contained a synaptic vesicle cluster labeled with synapsin and were associated with an elongated PSD-95 structure along the edge of the vesicle cluster as described previously ^20,39,52,72,74^. For intensity profile analyses, a ∼1000 nm long, 200 nm wide, rectangular ROI was drawn perpendicular and across the center of the PSD-95 structure, and the intensity profiles were obtained using this ROI for both the protein of interest and PSD-95. To align individual profiles, the PSD-95 signal only was smoothened using a rolling average of 5 pixels, and the smoothened signal was used to define the peak position of PSD-95. The profiles for the protein of interest (Ca_V_ or HA) and smoothened PSD-95 were aligned to the PSD-95 peak position, averaged across synapses, and then plotted. Peak intensities were also analyzed by extracting the maximal value from the line profiles of the protein of interest (Ca_V_ or HA) and smoothened PSD-95 within a 200 nm window around the PSD-95 peak. Peak intensity values were plotted for each synapse and averaged. For quantification of confocal images, a custom MATLAB program (https://github.com/hmslcl/3D_SIM_analysis_HMS_Kaeser-lab_CL) was used to generate masks of the presynaptic marker (synapsin), with the threshold determined by automatic two-dimensional segmentation (Otsu algorithm) ^75^. Regions of interest (ROIs) were defined as synapsin-positive areas formed by contiguous pixels of at least 0.05 μm^2^ in size. Each image typically contained between 500 and 1500 synapsin ROIs. Levels of HA or Ca_V_2.1 within these ROIs were measured and the average intensity across all ROIs within an image was calculated and plotted. Representative images in figures were cropped, rotated with bi-linear interpolation, and then brightness and contrast adjusted to facilitate inspection. Brightness and contrast adjustments were made for display in figures and were done identically for images within an experiment, but image quantification was performed on raw images without these adjustments. The experimenter was blind to the condition/genotype for image acquisition and analyses for STED and confocal microscopic experiments.

### Confocal imaging of neuronal somata

Neurons cultured on 0.17 mm thick 12 mm diameter (#1.5) coverslips were washed with PBS warmed to 37 °C and fixed in 2% PFA + 4% sucrose for 10 min at room temperature. Coverslips were then rinsed three times in PBS + 50 mM glycine at room temperature, permeabilized in TBP for 1 h at room temperature, and incubated in primary antibodies at for ∼48 h at 4 °C. The following primary antibodies were used: mouse IgG1 anti-HA (1:500, RRID: AB_2565006, A12) and mouse IgG2b anti-NeuN (1:500, RRID: AB_101711040, A254). After staining with primary antibodies, coverslips were rinsed twice and washed three times for 5 min in PBS + 50 mM glycine at room temperature. Alexa Fluor 555 (to detect HA; anti-mouse IgG1, RRID: 2535769, S19), and 633 (to detect neuronal somata; anti-mouse IgG2b, RRID: AB_1500899, S31) conjugated secondary antibodies were used at 1:500 dilution in TBP. Secondary antibody staining was carried out for ∼24 h at 4 °C. Coverslips were rinsed twice in PBS + 50 mM glycine, once in deionized water, air-dried and then mounted on glass slides using fluorescent mounting medium. Confocal images of neuronal somata were acquired on a Leica Stellaris 5 microscope with a 63x oil-immersion objective. Single section, 92.65 x 92.65 μm^2^ areas were acquired using 2x digital zoom (1024 x 1024 pixels, pixel size of 90.2 x 90.2 nm^2^). Imaging and laser settings were identical for all conditions within a given biological repeat. For analyses of somatic HA signals, the NeuN signal was used to mark the neuron somata, and EGFP-Cre or EGFP-ΔCre was used to define nuclei. Somatic ROIs were drawn as donut shapes by using the outer edge of the NeuN profile along the main somatic compartment not including neurites, and by excluding the EGFP-labeled nucleus. The average pixel intensity within the somatic ROI was then calculated for HA and plotted for each cell. Representative images in figures were cropped and adjusted for brightness and contrast to facilitate inspection. Brightness and contrast adjustments were made for display in figures and were done identically for images within an experiment, but image quantification was performed on raw images without these adjustments. The experimenter was blind to the condition/genotype for image acquisition and analyses.

### Electrophysiology

Electrophysiological recordings in cultured hippocampal neurons were performed as described previously ^20,39,74^ at DIV16 to 19. Glass pipettes were pulled at 2 to 5 MΩ and filled with intracellular solution containing (in mM) for EPSCs: 120 Cs-methanesulfonate, 2 MgCl2, 10 EGTA, 4 Na_2_-ATP, 1 Na-GTP, 4 QX314-Cl, 10 HEPES-CsOH (pH 7.4, ∼300 mOsm) and for IPSCs: 40 CsCl, 90 K-gluconate, 1.8 NaCl, 1.7 MgCl_2_, 3.5 KCl, 0.05 EGTA, 2 Mg-ATP, 0.4 Na_2_-GTP, 10 phosphocreatine, 4 QX314-Cl, 10 HEPES-CsOH (pH 7.2, ∼300 mOsm). Cells were held at +40 mV for NMDAR-EPSCs and at −70 mV for IPSCs. Access resistance was monitored during recordings and compensated to 2-3 MΩ, and cells were discarded if the uncompensated access exceeded 15 MΩ during the experiment. The extracellular solution contained (in mM): 140 NaCl, 5 KCl, 2 MgCl_2_, 1.5 CaCl_2_, 10 glucose, 10 HEPES-NaOH (pH 7.4, ∼300 mOsm), and recordings were performed at room temperature (20-24 °C). For NMDAR-EPSCs, picrotoxin (PTX, 50 μM) and 6-Cyano-7-nitroquinoxaline-2,3-dione (CNQX, 20 μM) were present in the extracellular solution. IPSCs were recorded in the presence of D-2-amino-5-phosphonopentanoic acid (D-AP5, 50 μM) and CNQX (20 μM) in the extracellular solution.

Action potentials were elicited with a bipolar focal stimulation electrode fabricated from nichrome wire. To evaluate the Ca_V_ blocker sensitivity of synaptic transmission, ω-agatoxin IVA (to block Ca_V_2.1) or isradipine (to block Ca_V_1s) were used. Blockers were pipetted into the recording chamber as concentrated stocks diluted in extracellular solution for a final working concentration of 200 nM for ω-agatoxin IVA and 20 µM for isradipine. For wash-in, cells were incubated after blocker addition for 5 min. IPSCs were recorded first in the absence of Ca_V_ blockers. Then, IPSCs were measured after wash-in of 200 nM ω-agatoxin IVA and again after wash-in of 200 nM ω-agatoxin IVA and 20 µM isradipine (Fig. 4F-I), or after wash-in of 20 µM isradipine (Fig. S5). Data were acquired at 5 kHz and lowpass filtered at 2 kHz with an Axon 700B Multiclamp amplifier and digitized with a Digidata 1440A digitizer. Data acquisition and analyses were done using pClamp10. For electrophysiological experiments, the experimenter was blind to the genotype throughout data acquisition and analyses.

### Western blotting

Lysates from transfected HEK293T cells were used for Western blotting. Ca_V_1 and Ca_V_2 constructs were co-transfected with Ca_V_β1b (p754; pMT2 Ca_V_β1b-GFP was a gift from A. Dolphin, Addgene plasmid # 89893; http://n2t.net/addgene:89893; RRID: Addgene_89893) ^76^ and Ca_V_α2δ1 (p752; CaVα2δ1 was a gift from D. Lipscombe, Addgene plasmid # 26575; http://n2t.net/addgene:26575; RRID: Addgene_26575) ^77^. Plasmids were transfected with the Ca^2+^ phosphate method at a 1:1:1 molar ratio with a total of 6.7 μg DNA. Around 48 h after transfection, HEK293T cells were harvested in 1 mL of standard 1x SDS buffer per flask. Homogenates were centrifuged at 16,200 x g for 10 min at room temperature, run on 6% (for Ca_V_s) or 12% (for β-actin) polyacrylamide gels, and transferred onto nitrocellulose membranes for 6.5 h at 4 °C in transfer buffer (containing per L, 200 mL methanol, 14 g glycine, 3 g Tris). Membranes were blocked in filtered 10% nonfat milk/5% goat serum in TBST (Tris-buffered saline with 0.1% Tween) for 1 h at room temperature and incubated with primary antibodies in 5% nonfat milk/2.5% goat serum in TBST overnight at 4 °C. The primary antibodies used were mouse IgG1 anti-HA (1:1000; RRID: AB_2565006, A12) and mouse IgG1 anti-β-actin (1:2000; RRID: AB_476692, A127). Membranes were washed five times for 3 min each at room temperature in TBST and then incubated with secondary antibodies in 5% nonfat milk/2.5% goat serum for 1 h at room temperature. The secondary antibodies used were peroxidase-conjugated goat anti-mouse IgG (1:10,000, RRID: AB_2334540, S52) and peroxidase-conjugated goat anti-rabbit IgG (1:10,000, RRID: AB_2334589, S53). Membranes were again washed five times for 3 min each at room temperature in TBST, then incubated in a chemiluminescent reagent for 30 s. Finally, the membranes were exposed to films, and films were developed and scanned. Corresponding western blots of Ca_V_s and β-actin were run simultaneously, on the same day, and on separate gels using the same samples. For illustration in figures, blots were rotated with bilinear interpolation and cropped for display.

### Quantification and statistical analyses

Data are displayed as mean ± SEM. Statistics were performed in GraphPad Prism 9, and significance is presented as *p < 0.05, **p < 0.01, and ***p < 0.001. Sample sizes and statistical tests for each experiment are included in each figure legend. For electrophysiological experiments, the sample size used for statistical analyses was the number of recorded cells. For STED microscopic data, the sample size used for statistical analyses was the number of synapses. For confocal microscopic data, the sample size used for statistical analyses was the number of images for analyses of synapsin ROIs, or the number of neurons for analyses of somata. Single factor, multiple group comparisons were conducted using Kruskal-Wallis tests followed by Dunn’s multiple comparisons post-hoc tests for proteins of interest (HA or Ca_V_2.1) and for current amplitudes (EPSCs, IPSCs). To compare the efficacy of blockade of synaptic transmission by different pharmacological agents in Fig. 4H, Friedman tests and Dunn’s multiple comparisons post-hoc tests were used. To compare the effects of Ca_V_ blockers on synaptic transmission across genotypes in Fig. 4I, two-way, repeated-measures ANOVA and Dunnett’s multiple comparisons post-hoc tests were used. In Fig. S5, the Wilcoxon matched-pairs signed rank test was used.

